# Neural connectivity of a computational map for fly flight control

**DOI:** 10.1101/2025.05.29.656834

**Authors:** Serene Dhawan, Zijin Huang, Bradley H. Dickerson

## Abstract

Nervous systems rely on sensory feature maps, where the tuning of neighboring neurons for some ethologically-relevant parameter varies systematically, to control behavior^1,2^. Such maps can be organized topographically or based on some computational principle. However, it is unclear how the central organization of a sensory system corresponds to the functional logic of the motor system. This problem is exemplified by insect flight, where sub-millisecond modifications in wing-steering muscle activity are necessary for stability and maneuverability. Although the muscles that control wing motion are anatomically and functionally stratified into distinct motor modules^3–7^, comparatively little is known about the architecture of the sensory circuits that regulate their precise firing times. Here, we leverage an existing volume of an adult female VNC of the fruit fly *Drosophila melanogaster*^8,9^ to reconstruct the complete population of afferents in the haltere–nature’s only biological “gyroscope”^10,11^–and their synaptic partners. We morphometrically classify these neurons into distinct subtypes and design split-GAL4 lines that help us determine the peripheral locations from which each subtype originates. We find that each subtype, rather than originating from the same anatomical location, is comprised of multiple regions on the haltere. We then trace the flow of rapid mechanosensory feedback from the peripheral haltere receptors to the central motor circuits that control wing kinematics. Our work demonstrates how a sensory system’s connectivity patterns construct a neural map that may facilitate rapid processing by the motor system.

---

To navigate complex environments, animals must rapidly detect and transform sensory information into precise motor commands. In flies, flight control relies on mechanosensory feedback from the haltere, a multifunctional gyroscopic organ derived from the hindwing^10–12^. Halteres are essential for flight; flies crash catastrophically without them^11,13^. Body rotations, resulting from either active maneuvers or passive external perturbations, create complex patterns of strain across the haltere’s surface that are encoded by specialized cuticular structures known as campaniform sensilla^10,11,14^. The haltere surface possesses dense arrays of campaniforms arranged in four highly stereotyped, spatially-distinct fields^10,15–19^. A wealth of behavioral evidence implicates the halteres in a range of stability reflexes via subtle adjustments in head and wing kinematics that require sub-millisecond control^14,20–25^. However, previous anatomical and electrophysiological work has only provided a rough picture of the neural circuit architecture. The haltere campaniform afferents send their axonal projections into the ventral nerve cord (VNC), the fly analog of the vertebrate spinal cord^12,18,26^, and brain. Yet, beyond the connection to a single wing steering muscle motor neuron, the specific synaptic targets of the haltere afferents have not been determined^18,27^. Moreover, the campaniform sensilla are arranged into four distinct fields on the haltere itself, but it is unknown whether this organization is preserved in downstream circuits or whether the haltere afferent projections are re-mapped based on their role in flight control. The latter organization might allow motor networks to more efficiently integrate haltere feedback, but has not yet been demonstrated. Here we harness the power of connectomics to describe the connectivity of the haltere afferents and reveal a map for the control of wing motion.

## Morphology-based cell typing

In *Drosophila*, sensory afferents enter the VNC through a highly stereotyped set of peripheral nerves, making it possible to assign a sensory modality to some axons based solely on their gross anatomy^28^. Guided by the knowledge that the axonal projections of the haltere afferents enter the VNC through the dorsal metathoracic nerve, we identified this structure in the FANC electron microscopy dataset^8,9^ and subsequently reconstructed all neurons within it. This resulted in 372 candidate haltere afferents (Fig. 1A) whose identity we validated against light microscopy images of established genetic driver lines^29^.

**Figure 1:**
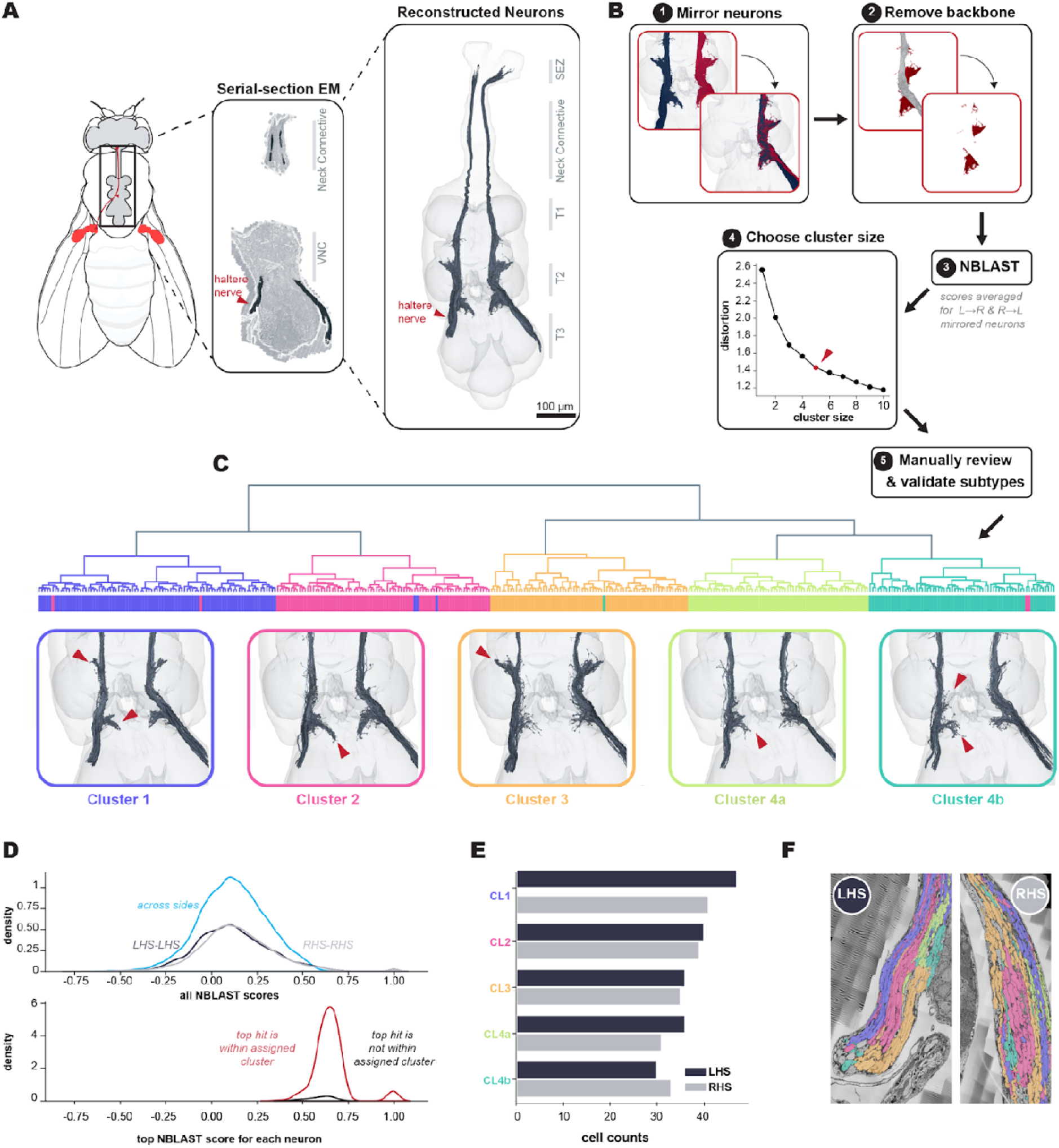
Reconstruction & cell-type classification of the haltere primary sensory afferents. **A:** Schematic of an adult fly with the halteres and their associated sensory afferents depicted in red. A serial-section electron micrograph, taken from a volume of an adult female ventral nerve cord (VNC), highlights the nerve tract through which these sensory afferents enter the VNC. The fully reconstructed set of sensory afferents (*n = 372*), found by sampling this tract, are shown at the right. **B:** Pipeline for classifying the haltere sensory afferents into morphological subtypes. Neurons are mirrored onto a common side, followed by manual isolation of the terminal arbors. Pairwise morphological similarity scores are calculated for these segments using NBLAST and the final scoring matrix is determined by taking the mean of the scores for left-to-right vs right-to-left mirrored neurons. Scores are hierarchically clustered and the final group size is determined using the ‘elbow method’. Neurons within each cluster are subjected to manual review. **C:** Dendrogram showing morphological clustering of haltere sensory afferents with coloured branches and leaf nodes denoting different neuronal subtypes. Renderings of each subtype are shown with arrows highlighting their key anatomical features. **D:** The distribution of NBLAST scores for pairwise morphological comparisons between neurons that originate on the same side of the VNC vs different sides is largely the same (*top panel*). The top-scoring, non-self match for each neuron is generally within its assigned morphological cluster (*bottom panel*). **E:** Left vs right cell counts for each morphological subtype of haltere afferent. **F:** Example serial-section EM slice of the left and right haltere nerve tracts, with neurons colored by morphological subtype.

We then adopted a morphology-based approach and classified the complete population of haltere afferents into distinct subtypes. This enabled a more efficient interpretation of their synaptic connectivity, by effectively ‘compressing’ multiple cells in the network into subtypes. To this end, we mirrored the axonal terminals of the haltere afferents onto a common hemisphere and generated pairwise similarity scores using NBLAST^30^ (Fig. 1B). Next, we hierarchically clustered our NBLAST scores and assigned neurons to one of five groups (Fig. 1C). We followed this with an extensive manual review of within- and across-subtype stereotypy in fine branching structure and the reassignment of 12 neurons. Notably, the distribution of scores pertaining to within vs across-side comparisons were indistinguishable (Fig. 1D). This suggests that there is a low degree of biological variability in neuronal morphologies across hemispheres and that inaccuracies in the mirroring transformation did not significantly impact scoring (Fig. 1D).

Our finalized subtypes produce a balanced number of neurons within each cluster across hemispheres (Fig. 1E), and importantly, in >99% of cases, the top-scoring NBLAST match for each neuron corresponds to another member of its own subtype (Fig. 1E). Intriguingly, we observed that neurons within a subtype appeared to fasciculate and enter the VNC as distinct bundles within the dorsal metathoracic nerve (Fig. 1F).

## Diverse peripheral origins of subtypes

Our reconstruction of the haltere afferents suggests that there is a clear pattern regarding how haltere mechanosensory axons project into the central nervous system. However, the FANC dataset does not include the full peripheral nerves of the body, making it unclear how these morphological subtypes map onto the haltere campaniform fields. Moreover, the encoding properties of each campaniform are likely determined by its location and local mechanics during oscillation^31^. Determining the origins of each morphological cluster would therefore provide a more nuanced understanding of how the five haltere afferent subtypes might differentially regulate flight control. Additionally, it would also test whether haltere feedback has a topographic representation in the VNC or if haltere input is organized based on some behaviorally relevant parameter.

We matched each subtype against light microscopy databases^32^ to construct split-GAL4 lines and imaged GFP expression in the VNC and halteres of individual flies (Fig. 2B); cluster 4b exists as a publicly available driver line^32^. Our imaging indicates that the haltere projections into the VNC cannot be captured by a simple topographic mapping of their arrangement at the periphery (Fig. 2C-D). That is, each haltere afferent subclass comprises neurons from multiple campaniform fields rather than consisting solely of neurons from a single campaniform field (Fig. 2E).

**Figure 2:**
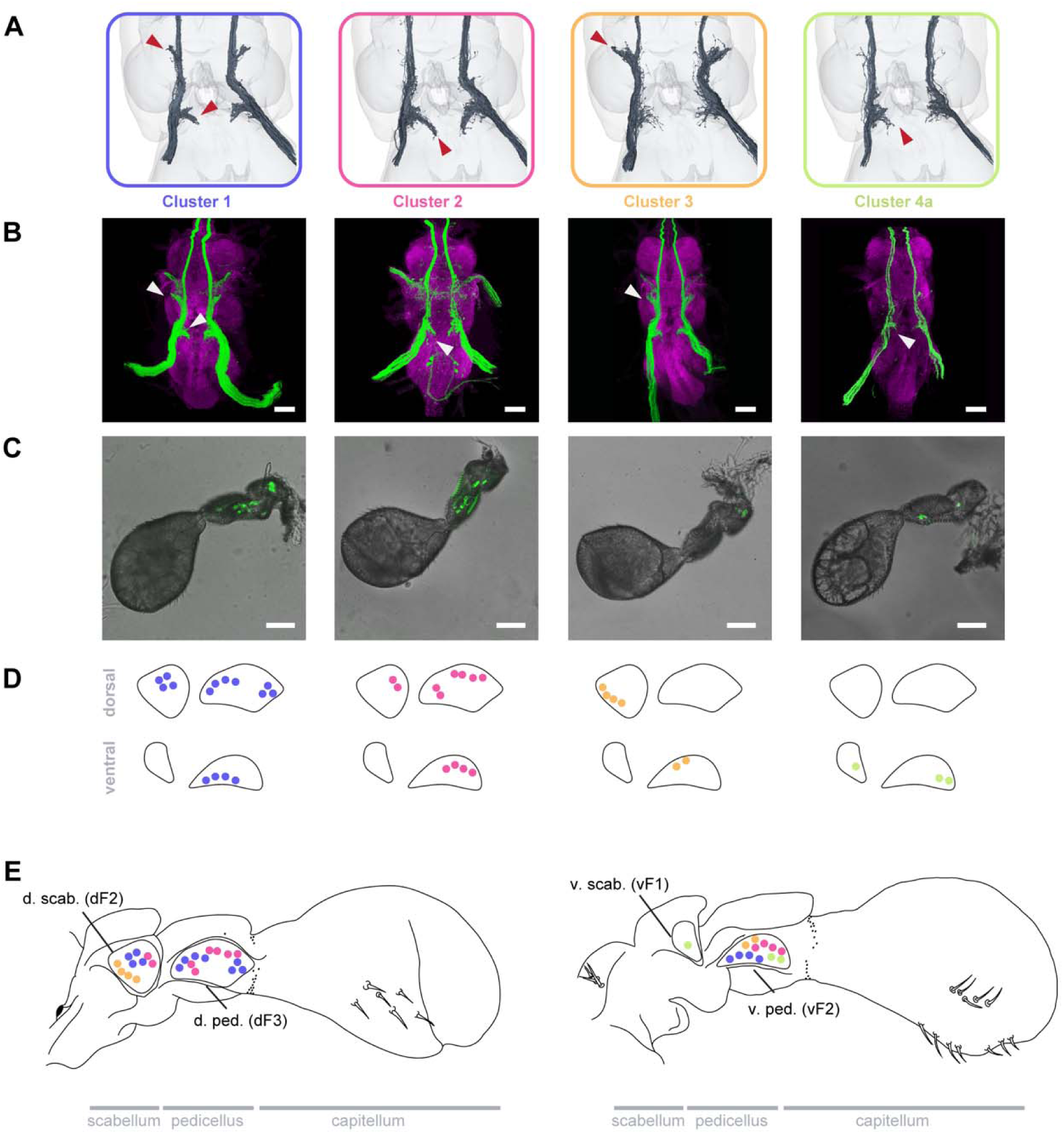
Peripheral origins of haltere afferent subtypes. **A**: EM reconstructions of the haltere afferents. **B:** Maximum intensity projections of the VNC showing GFP expression driven by split-GAL4 lines which separately label each haltere afferent morphological subtype. Scale bars: 50µm **C:** Corresponding light micrographs for each split-GAL4 line showing GFP expression on the peripheral haltere campaniform fields. Scale bars: 50µm **D:** Schematic of the dorsal and ventral haltere campaniform fields, with colored dots representing the peripheral locations of each afferent subtype. **E:** Schematic of the *Drosophila* haltere, showing the anatomy of the four major campaniform fields (adapted from Cole and Palka^12^). Haltere afferent subtype locations are overlaid from D.

## Output connectivity of haltere afferents

Having mapped the peripheral origins of the haltere afferents, we next sought to characterize the central circuitry integrating haltere mechanosensory information. We began by quantifying the number, type, and spatial distribution of their synaptic sites, which are automatically predicted. Across subtypes, our EM reconstructions indicate that the haltere afferents possess a large number of synaptic outputs (∼400/neuron) with significantly fewer inputs (∼15/neuron) by comparison (Fig. 3A). The majority of these inputs show a pattern of axo-axonic connectivity between haltere afferents within a subtype (Extended Data Fig. 1A). Reciprocal connections between excitatory ‘sister’ neurons have previously been reported in both the *Drosophila* and vertebrate olfactory systems^33^, and may increase signal robustness through correlated transmitter release^34–36^.

**Figure 3:**
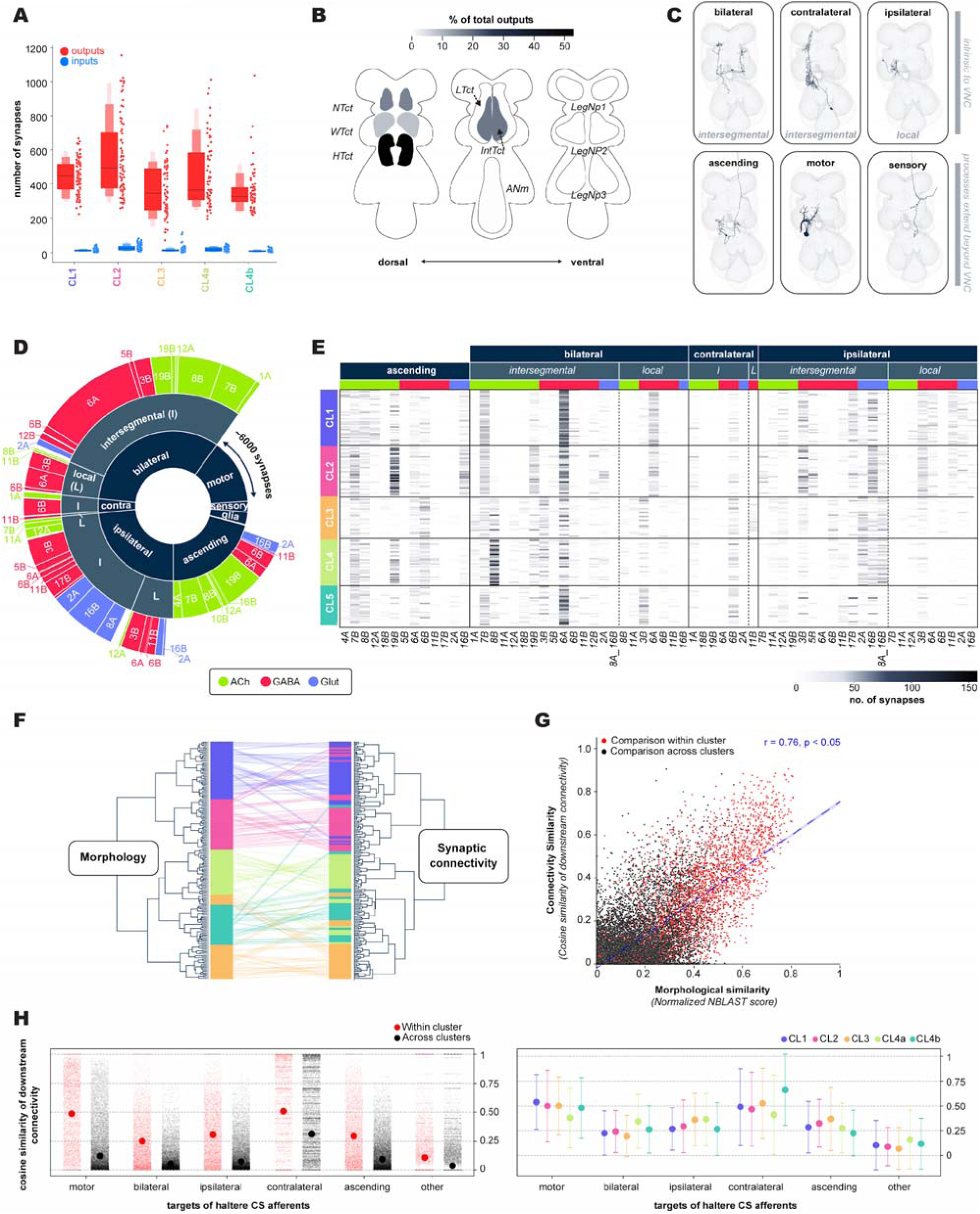
Reconstruction of the synaptic outputs of the haltere afferents reveals subtype-specific connectivity. **A**: Quantification of the total number of synaptic inputs (blue) and outputs (red) for each morphological subtype of haltere afferent. **B:** Heatmap depicting the spatial distribution of the haltere synaptic outputs within the different neuropils of the VNC. The upper tectulum consists of the neck, wing and haltere tectulums *(NTct, WTct, HTct*). Other major neuropils include the intermediate tectulum (*IntTct*), lower tectulum (*LTct*), abdominal neuromere (*ANm*) and the leg neuropils (*LegNP1-3*). **C:** Example renderings of different broad classes of neuron that receive input from the haltere afferents. These can be grouped into neurons intrinsic to the VNC – which are further subdivided into cell types that are restricted to one neuropil (local) vs types that project across neuropils (intersegmental) – and neurons that project out of the VNC to either the body or central brain. **D:** Sunburst chart showing the breakdown of the targets of the LHS haltere sensory afferents. The innermost layer is segmented into broad classes of neuron, the middle layer subdivides these classes into local vs intersegmental where applicable, and the outermost layer further classifies neurons by their developmental hemilineage and associated transmitter identity (acetylcholine = green, GABA = red, glutamate = blue). **E:** Heatmap showing outputs from the LHS haltere afferents to their target neurons. Rows are colored by haltere morphological subtype; columns are grouped by broad cell class and then subdivided by projection pattern and hemilineage. **F:** Tanglegram comparing hierarchical clustering of pairwise morphological similarity scores (NBLAST scores) vs connectivity similarity scores (cosine similarity of downstream synaptic connectivity). Individual neurons are connected by colored inter-tree edges. **G:** For each pairwise neuronal comparison of the haltere sensory afferents, morphological similarity is plotted against connectivity similarity. Comparisons within a haltere afferent subtype are depicted in red whereas comparisons across subtypes are shown in black. **H:** The cosine similarity of the synaptic outputs onto different broad classes of neuron from haltere afferents within vs across subtypes (*left panel)* and within-haltere-subtype similarity of downstream connectivity onto different broad classes of neuron (*right panel*).

In accordance with previous work implicating the haltere in both flight control and gaze stabilization^22,24,25^, the synaptic outputs of the haltere afferents are primarily localized to the upper tectulum (Fig. 3B), which contains the circuitry underlying these behaviors^28^. The remainder of the haltere outputs are distributed within the intermediate tectulum, an integrative area hypothesized to coordinate leg and wing motion^37^. This finding putatively suggests a novel function for the haltere in coordinating limb motion during take-off and grooming. The patterns of the output synapses are largely invariant across haltere afferent subtypes (Extended Data Fig. 1B-C) and are recapitulated by the spatial distribution of haltere inputs (Extended Data Fig. 1D)

To obtain a more fine-grained understanding of the circuitry responsible for processing haltere mechanosensory information, we reconstructed all postsynaptic neurons predicted to receive at least 4 synapses from the haltere afferents. We focused our proofreading efforts on the left side of the VNC as it is more intact than the right^38^. The result is 4479 neurons that receive 69199 synapses from the haltere afferents (Extended Data Fig. 1E). To systematically organize these neurons, we defined a hierarchical set of annotations (Fig. 3C-D). First, we assigned all reconstructed outputs of the haltere afferents to a broad class (see Methods for full details). We defined bilateral, contralateral and ipsilateral neurons as being intrinsic to the VNC, then further classified neurons as local or intersegmental depending on whether their projections spanned multiple neuromeres^28^ Neurons with processes extending beyond the VNC included ascending and descending neurons, other sensory afferents and motor neurons. For our next level of annotations, we assigned neurons to a hemilineage, i.e., a group of neurons that arise from a single neuroblast during development and that share a neurotransmitter identity^39,40^ (Fig. 3E, Extended Data Fig. 1F, Extended Data Fig. 2).

To address whether morphology-based classification of the haltere afferents recovers functionally-relevant groupings, we quantified the similarity of their downstream connectivity using the cosine similarity metric. Afferents which connect to similar targets in similar proportions; i.e., those with high cosine similarity, are likely to play similar roles in information processing and behavior. Supporting the idea that our morphological subtypes reflect functional units, hierarchical clustering comparing morphological versus synaptic connectivity reveals that the major subtypes are largely preserved across clustering methods whereas the ‘within-subtype’ orders are somewhat shifted (Fig. 3F). Further, we find that connectivity-based measures of neuronal similarity correlate well with morphology-based similarity measures (Fig. 3G).

We next tested whether the haltere afferents exhibit subtype-specific connectivity with all broad classes of neuronal targets or whether the high within-subtype cosine similarity scores are biased to a single class of target neuron. To answer this, we calculated separate cosine similarity scores for the connectivity vectors of the haltere afferents with their targets grouped by broad subtype (Fig. 3H). We find that within-subtype connectivity similarity is highest in relation to motor neurons and contralateral neurons.

## Afferent subtypes target motor modules

Ultimately, mechanosensory feedback from the haltere dynamically structures the timing and activation of the wing-steering system. The haltere afferents provide strong excitatory input via monosynaptic connections to the wing steering muscle motor neurons (MNs). Each steering muscle is innervated by a single motor neuron, which have all been uniquely identified in the FANC connectome^8,38^. The activity of the steering muscles changes the conformation of the wing hinge, resulting in subtle changes in wing kinematics^41–43^. These muscles can be classified anatomically based on the specific cuticular element, or sclerite, to which they attach. The steering muscles attach to four sclerites, each of which is hypothesized to control specific aspects of wing motion: the basalares; the first axillary; the third axillary; and the fourth axillary (Hg). The steering muscles are also functionally stratified as ‘tonic’ or ‘phasic’. Tonic muscles mediate continuous, fine-tuned motor control whereas phasic muscles execute large, transient changes in wing motion during active maneuvers (Fig. 4A)^3,5,6^.

**Figure 4:**
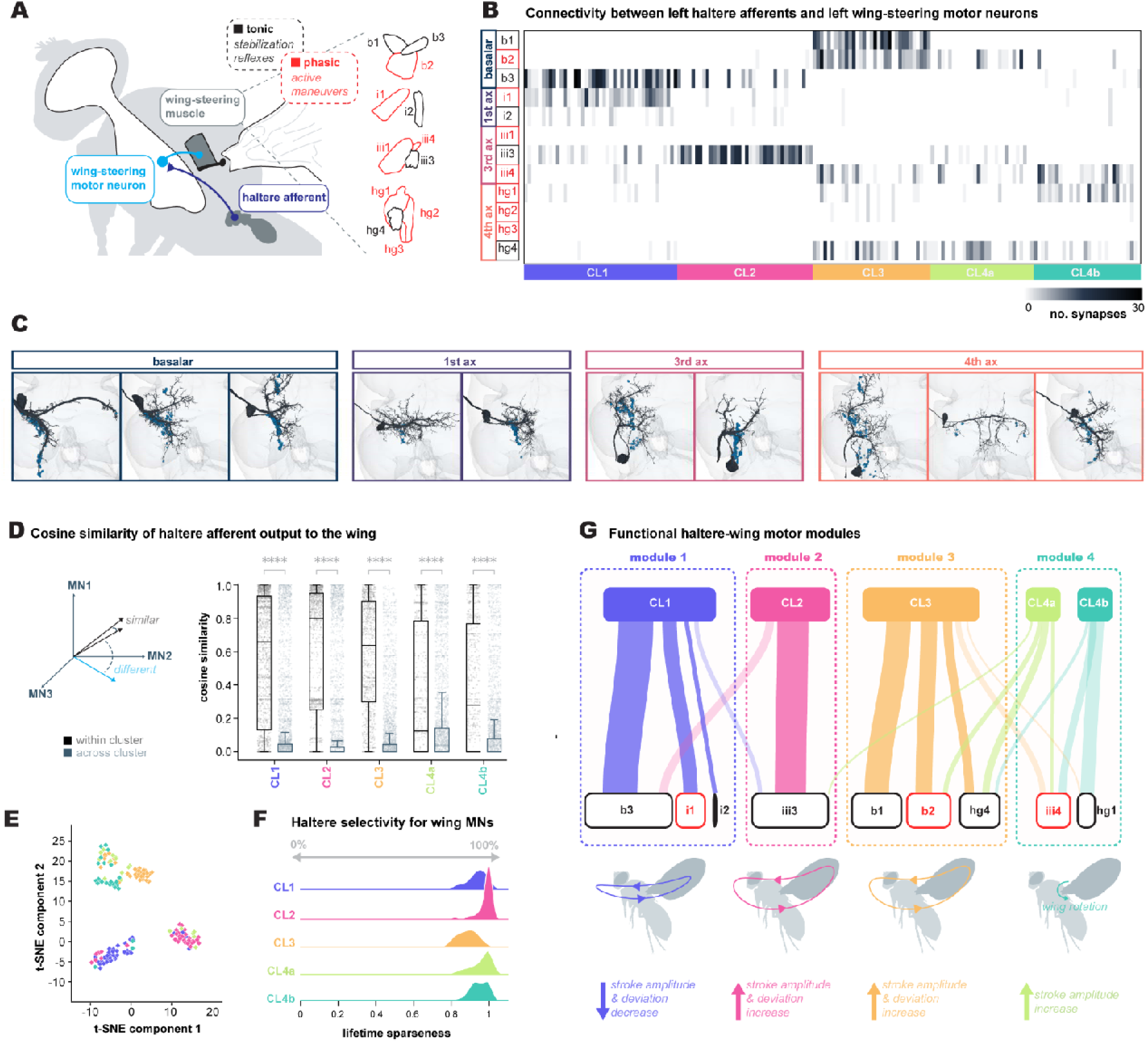
Haltere morphological subtypes preferentially target motor modules with a common behavioral output. **A**: Schematic showing the anatomy of the *Drosophila* wing steering system. A wing steering muscle motor neuron innervates a single muscle which in turn attaches onto a specific cuticular element or sclerite on the wing hinge. The muscles associated with each sclerite can be further subdivided into two functional classes: muscles that are tonically active (black) or phasically recruited (red) during flight. Synaptic outputs from a haltere sensory afferent onto a wing-steering motor neuron in part determine the motor neuron’s spike-timing, thereby altering the movement of the wing. **B:** Connectivity between the LHS haltere sensory afferents and the wing-steering motor neurons. Rows are grouped by the sclerite which a wing steering muscle attaches and colored by tonic (black) or phasic (red) activity patterns of the associated muscle; columns are colored by haltere morphological subtype. **C:** Renderings of the wing steering muscle motor neurons with synaptic inputs from the haltere sensory afferents depicted in blue. **D:** Cosine similarity of vectors for synaptic connectivity between the haltere afferents and the wing steering muscle motor neurons, where each value in the vector is the normalized number of synapses from a single haltere afferent to a single motor neuron. Connectivity similarity of neurons is higher within a haltere afferent morphological subtype vs across subtypes (ANOVA; Conover’s post hoc pairwise test with Holm correction for multiple comparisons ns, not significant; *p < 0.05; **p < 0.01; ****p < 0.0001). **E:** t-SNE plot of synaptic connectivity vectors between the haltere afferents and wing-steering motor neurons. **F:** Lifetime sparseness (LTS) of the haltere afferent output to the wing. Across all morphological subtypes, the distribution of LTS values is close to 1. **G:** Summary wiring diagram of connectivity between the haltere afferents and wing steering muscle motor neurons. Each class of haltere morphological subtype directly synapses onto a distinct and largely non-overlapping subset of wing steering muscle motor neurons that share a common behavioral function.

To date, only a single example of direct connectivity between the haltere and wing has been confirmed via anatomical and electrophysiological work in reduced preparations^18,27^. Our analysis identified extensive synaptic connectivity between the haltere afferents and wing-steering MNs (Fig. 4B-C). Given widespread damage within the sensory domain of the right wing nerve of FANC, we concentrated our analysis on the left wing MNs. Although raw synapse numbers are reduced on the right it is notable that the connectivity patterns identified here are consistent across hemispheres (Extended Data Fig. 3A-B), thus indicating that our findings are not an idiosyncratic feature of a limited portion of the FANC dataset.

A qualitative examination of the connectivity matrix indicates that the haltere afferents exhibit subtype-specific connectivity to distinct subsets of wing-steering MNs (Fig. 4B). To validate this observation, we computed cosine similarity scores for the normalized output vectors of each haltere afferent-wing MN pair. As expected, within-subtype comparisons yield higher cosine similarity values than across-subtype comparisons. Values for across-subtype comparisons are distributed close to 0, suggesting that there is almost no overlap in connectivity patterns between different haltere afferent subtypes (Fig. 4D). Indeed, dimensionality reduction techniques capture the unique output ‘fingerprint’ of each subtype (Fig. 4E). To quantify the degree of selectivity for each haltere afferent subtype with respect to the wing-steering MNs it targets, we calculated lifetime sparseness (LTS) scores^44^. In this context, an LTS of 0 is indicative of a haltere afferent which connects to all 12 wing-steering MNs in equal proportions, whereas an LTS of 1 is indicative of an afferent that connects to only 1 wing-steering MN. Across all haltere afferent subtypes, LTS scores are close to 1 (Fig. 4F).

## Discussion

The canonical model of haltere function holds that field dF2 is maximally responsive to the shear strains resulting from gyroscopic forces whereas dF3 detects in-plane inertial strains as the haltere oscillates^10,11^. Accordingly, previous investigations of haltere function have treated afferents originating from the same campaniform field as a discrete functional unit^27,45^. For example, previous work in the blowfly *Calliphora* identified a monosynaptic pathway between a substantial fraction of neurons from the second dorsal field (dF2) and the b1 wing steering muscle motor neuron that is proposed to mediate stabilization reflexes^18,27^. In *Drosophila*, the b1 motor neuron also receives haltere input^26^, yet the peripheral origin of this feedback was unknown. The broad connectivity between the haltere afferents and the wing steering muscle motor neurons that we describe suggests that the haltere recruits specific combinations of muscles to correct mechanical perturbations.

Although our work highlights the importance of the haltere in mediating flight control, our reconstruction efforts only provide a static picture of the haltere circuitry. Still, these results are consistent with the emerging view that each campaniform field is continuously active during flight and helps regulate the entire wing steering network, even in the absence of body rotations^29,46^. Indeed, recent work implicates the haltere campaniform fields in regulating both stabilizing and active flight maneuvers via the functional segregation of the haltere steering muscles^45^ that receive descending visual input^29,47^. We found that haltere afferents within four of the five identified subtypes target both tonically and phasically active muscles. Thus, the circuitry of each campaniform subtype predicts that haltere input helps the wing steering system achieve its high dynamic range necessary for flight maneuvers. Precisely how the different classes of haltere afferents are individually regulated to bias the activity of the wing steering muscles during flight remains unclear, especially as campaniforms are sensitive to nanometer-scale deflections^48^.

### Functional implications for flight

The haltere’s campaniform fields not only synapse onto the wing steering muscle motor neurons of both functional types, these projection patterns form putative motor modules. The motor modules for wing motion that are determined by all premotor neurons synapsing onto the wing steering muscle motor neurons are mostly anatomically clustered based on the sclerite to which each muscle attaches^38^. By contrast, here we found that the modules formed by each haltere afferent subtype consist of muscles attached to different sclerites that likely act synergistically on wing motion (Fig. 4G). This is consistent with recent machine learning approaches to understanding the mechanics of the wing hinge, which demonstrate that there are complex relationships among the steering muscles across different sclerite groups^48^. For example, CL1 afferents target b3, i1,and i2. The activity of b3 and i1 are strongly correlated with decreases in amplitude and deviation^4–6,49^. By contrast, CL2 afferents almost exclusively target iii3. Previous epifluorescent calcium imaging of the haltere campaniforms indicates that their activity is correlated with wingstroke amplitude^45^. Similar imaging of the wing steering muscles shows that iii3 is the wing steering muscle whose activity is most strongly correlated with this parameter^6^. Past dye fills of the wing steering motor neurons suggest that the morphology of iii3 may facilitate precise, high-frequency firing via chemical and electrotonic synapses^50^. Although electrical synapses currently cannot be resolved with EM, the surface area overlap between CL2 afferents and iii3 suggests that their existence is a strong possibility (Extended Data Fig. 3C-E). Looking across the wing steering system, the putative electrical synapses we identified recapitulate the subtype-specific connectivity matrix using chemical synapse prediction (Extended Data Fig. 3C-E). This suggests that a mix of chemical and electrical communication is a general property of flight control circuitry. For example, anatomical and physiological evidence confirms that haltere afferents supply the b1 motor neuron with both chemical and electrotonic input^18,26,27,51^. Moreover, b1–like iii3–exhibits a distinctive ‘stubbly’ dendrite morphology, a large axon, and fires each wingstroke^3–5,52^. The b1 motor neuron receives input from CL3 haltere afferents, as do the b2 and hg4 motor neurons. Previous experimental evidence demonstrates that b1 and b2 are responsible for increases in wingstroke amplitude and deviation^3,5,49,53^, whereas the hgs are hypothesized to regulate these same aspects of wing motion as well as the wing’s angle of attack^6,42,49^. Finally, CL4a and CL4b neurons have similar connectivity profiles and connect to muscles that may control wing camber and angle of attack^6,42,49^. The genetic reagents we developed will help uncover how each morphological subtype mediates specific flight maneuvers. Intriguingly, even when including disynaptic connections, haltere subtype-specific connectivity appears to be preserved within wing premotor networks (Extended Data Fig. 4A-B). Preferential connectivity between haltere subtypes and large, putatively electrically coupled interneurons^26,54^ may facilitate the bilateral coordination of wing movements (Extended Data Fig. 4C-D”). Crucially, all of the identified modules are based on the reorganization of the sensory periphery as the haltere afferents enter the VNC.

### A neural map of haltere information

The complex patterning of the terminal arborizations of the haltere afferents cannot be recapitulated by point-to-point topographic projections from the sensory epithelium, as found in many other systems^55–58^. Instead, afferents originating from multiple campaniform fields converge as they enter the VNC and form morphologically-distinct subtypes that share synaptic targets and putative functional roles (Fig. 2E). Past physiological characterizations of campaniform sensilla indicate that, in the context of locomotion, the encoding properties of these neurons is largely determined by the sensor’s location and resulting local biomechanics^31^. In addition, when subject to periodic oscillations, the neurons within campaniforms fire single action potentials at different preferred firing phases^59–62^. Our finding that each morphological subtype contains afferents from multiple fields suggests that each cluster provides feedback by sampling different preferred phases of the haltere stroke. Because of the varied peripheral origins of each morphological cluster, we predict that the neurons of a given subtype do not all fire at the same preferred phase. Rather, each subtype encodes distinct points in the haltere stroke, subsampling the haltere population code. Both types of steering muscles fire at precise times in the stroke cycle; we hypothesize that the peripheral locations for a given haltere afferent subtype help bias the firing phase of its steering muscle motor neuron targets.

Although it is not yet possible to determine the precise phase range of each subtype, our connectivity matrix combined with the genetic reagents we generated make testable predictions. For example, the motor neurons for b1 and b2 both fire near the upstroke-to-downstroke transition, and receive input from a subtype of afferents that primarily originate from dF2 (Fig. 2)^3,4,27^. During flight, the haltere beats antiphase to the wing, suggesting that dF2–which is positioned where in-plane strain during flapping is greatest–is maximally active during the predicted firing times of b1 and b2. Furthermore, the haltere helps mediate gaze stabilization via input to the neck motor system^20,25^; our results suggest the potential for a parallel map that indirectly regulates visual input. We anticipate that these new genetic resources will help reveal how population coding from each morphological subtype structures the firing phase of the wing steering muscle motor neurons and flight posture.

More broadly, the central organization of haltere feedback mirrors the organization of sensory input to facilitate efficient processing for rapid control found in other systems. In crickets, mechanosensory afferents with similar tuning profiles, as opposed to those that innervate adjacent filiform hairs, overlap extensively within the CNS and form an orderly representation of air current direction^63,64^. Such organization has also been described in vertebrates^1^, with perhaps the best characterized example being the auditory map of amplitude spectrum and time interval found in the midbrain of the barn owl^65,66^. An important distinction in the case of the haltere is that its afferents, in addition to communicating with a large population of interneurons for further processing, directly interface with the motor system. This emphasizes the critical role the haltere plays in regulating the biomechanics of the wing steering muscles via sub-millisecond precision spike timing.

### Conclusions

As dipterans radiated, the haltere campaniform fields hypertrophied, forming dense arrays whose connectivity patterns with the wing steering system are hypothesized to facilitate flies’ aerial abilities^10,11,18,19^. Indeed, the arrangement of the haltere afferents reflects the computational needs of wing motor circuits during flight. The existence of a connectivity map of haltere mechanosensory information invites the question of what developmental processes govern the formation of this circuit. In the case of the wing campaniform afferents, the choice of central pathway correlates directly with time of neuronal birth and physiological adaptation rates as opposed to the topographic distribution of receptors^67,68^. Moreover, individual campaniforms from a single field exhibit similarly diverse projection patterns as the haltere^69^. Given the serial homology between the haltere and wing, it is likely that developmental timing plays a significant role in generating neuronal diversity among the haltere afferents. With multiple emerging VNC connectomes^8,70^, future investigations exploring how the haltere’s wiring logic is developmentally controlled and varies across dipterans will provide powerful insight into the evolution of one of life’s most successful orders^71^.

## Methods

### Connectomic reconstruction within the FANC dataset

Previous work has described machine learning tools for automated neuron segmentation and synapse prediction^8^ within FANC (Female Adult Nerve Cord)^8^, a serial section transmission electron microscopy dataset containing the VNC, neck connective, and a limited portion of the brain’s subesophageal ganglion. We used Google’s collaborative Neuroglancer interface^72^ to manually correct errors in the automated segmentation of neurons of interest.

### Identification and reconstruction of the haltere campaniform afferents

The axons of the haltere campaniform afferents enter the VNC through the dorsal metathoracic nerve^28^. We chose multiple cross sections through the base of this nerve tract on both the left and right sides of the VNC and annotated every profile within these cross sections. We reconstructed each profile to ‘identification’, here defined as completion of the neuron’s large, microtubule containing backbone, and then matched our reconstructions against existing light microscopy images of the haltere nerve^29^. We then proofread candidate haltere campaniform afferents to ‘completion’, i.e. reconstruction of the entire neuronal arbor.

### Reconstruction of downstream synaptic partners of the haltere afferents

We used the automated synapse prediction to identify all neuronal objects which receive input from the haltere afferents (n = 144,157). We removed synapses with an associated confidence score below 20 to reduce the impact of spurious connections. Due to rough dissection of the right-hand side of the VNC^8^ we focused further reconstruction and identification efforts on the left hemisphere (n = 71,338). We proofread all neuronal objects which received at least 4 haltere inputs (n = 4479) until they were a) associated with a completed backbone with a cell body attached or b) identifiable as part of a known sensory or c) descending process. We categorized the remaining objects as fragments. A portion of neurons downstream to the haltere afferents (n=473) – including the wing motor neurons^8,38^ as well as several ascending and descending neurons^73^ – had been already proofread as part of independent studies. We accessed annotations for these neurons through the Connectome Annotation Versioning Engine (CAVE)^74^.

### Cell-typing analyses

#### Morphology-based classification of the haltere afferents

We downloaded haltere afferent meshes and then skeletonized these using the wavefront method implemented in the skeletor package (https://github.com/navis-org/skeletor). We manually removed neuronal backbones, i.e. the long straight portions of a neurite extending from the metathoracic neuromere towards the cervical connective, as their fasciculation reduced the sensitivity of the NBLAST algorithm^30^ to differences in axonal branch morphology. We scaled the resulting skeletons from units of nanometers to microns and resampled these to 1μm step size prior to conversion to the vector cloud dotprops format. We transformed the dotprops to be on the same side of the VNC, by mirroring those from afferents originating on the left side to the right, or vice versa, using the flybrains package (https://github.com/navis-org/navis-flybrains).

We calculated pairwise morphological similarity scores using NBLAST^30^ as implemented in the navis package (https://github.com/navis-org/navis). We calculated the final scoring matrix using the average of the NBLAST scores for left to right vs right to left mirrored afferents to account for errors in the mirroring transformation. We performed unsupervised hierarchical clustering of the scoring matrix, employing Ward’s clustering criterion, using scipy’s clustering package. The elbow heuristic, whereby distortion is plotted as a function of the number of clusters, indicated an optimal cluster number of 5. We validated this cluster size against three additional criteria: a) there were clear morphological differences between afferents in each cluster b) additional morphological differences were not uncovered by increasing the number of clusters and c) the number of neurons from each side of the VNC within each cluster was approximately equal. Following a determination of the total number of clusters, we manually reviewed the fine branching structure of each neuron and in a limited number of cases (n=12), we reassigned the neuron to a new cluster.

#### Morphology-based classification of the downstream synaptic partners of the haltere afferents

We used careful visual inspection to assign neurons to one of the following broad classes based on soma position and arborization patterns:

□ **Bilateral** neurons arborize on both sides of the VNC
□ **Contralatera**l neurons arborize solely on the opposite side of the VNC with respect to their soma position.
□ **Ipsilateral** neurons arborize on the same side of the VNC as their soma.
□ **Ascending** neurons have a soma in the VNC and send a process through the neck connective towards the central brain.
□ **Descending** neurons have no soma in the VNC and have a process through the neck connective
□ **Sensory** neurons enter the VNC via a peripheral nerve tract and have no soma in the VNC.
□ **Motor** neurons have a soma in the VNC and exit through a peripheral nerve tract.

The VNC can be divided into three bilaterally symmetrical thoracic neuromeres and one abdominal neuromere. We further classified intrinsic VNC neurons, i.e. the bilateral, contralateral and ipsilateral groups, as ‘local’, if their arborization was restricted to a single neuromere, or ‘intersegmental’ if their arbor spanned two or more neuromeres.

#### Neuronal hemilineage assignments

During VNC development, fixed sets of stem cells generate neurons in stereotyped hemilineages that share anatomical features, as well as a neurotransmitter identity ^39,75^. Notably, the primary neurite – i.e. the process extending directly from the cell body and which separates the axon from the dendrite – of neurons within a hemilineage converge to enter the neuropil as a cohesive bundle. With this in mind, we first grouped the downstream synaptic partners of the haltere afferents based on primary neurite location. We then manually matched these groups to known hemilineages based on light microscopy images of sparse GAL4 driver lines^40,75^ and neuroblast clones^75^, as well as previous annotations from a complete connectome of a male VNC^70^. Additional features used to aid identification included relative soma position and fine dendritic/axonal branching morphology. Finally, we assigned neurotransmitter identity by referencing work which has systematically mapped neurotransmitter choice across all VNC hemilineages^39^.

### Connectivity analyses

#### Raw synapse counts

We accessed automated synapse predictions through CAVE^74^ and set a threshold based on an associated confidence score of >20 connections prior to further analysis.

#### Quantification of the spatial distribution of synapses

We obtained VNC neuropil region meshes generated from a segmentation of the adult VNC^28^, which had been transformed into the FANC coordinate space, from the FANC GitHub (https://github.com/htem/FANC_auto_recon/tree/main/data/volume_meshes). To calculate whether the location of a synapse was within a particular neuropil mesh we used Ncollpyde (https://pypi.org/project/ncollpyde/).

#### Cosine similarity

Cosine similarity provides a measure of whether a given pair of neurons connect to the same targets in the same proportions. If the two neurons have no shared synaptic partners, their cosine similarity will be 0. Conversely, if these neurons target the exact same set of partners with the exact same number of synapses, their cosine similarity will be 1. We calculated cosine similarity scores using the cosine similarity method from scikit-learn.

#### Lifetime sparseness

Lifetime sparseness (LTS)^44^ provides a measure for the selectivity of a neuron’s connectivity pattern, where *N* is the total number of observations and *r_j_*is an observation’s value:

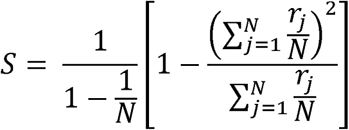

The higher the LTS value, the more selective the neuron. As pertains to the haltere afferents, an LTS of 0 would indicate an afferent that provides an equal number of synapses to all 12 of the wing-steering MNs whereas an LTS of 1 would indicate an afferent that synapses onto only 1 wing-steering MN. We performed LTS calculations using the implementation in the navis package (https://github.com/navis-org/navis).

#### Putative electrical synapses

Monosynaptic electrical connections have previously been described between the haltere afferents and the wing-steering MNs^27^. However, the size of electrical synapses (10-20 nm)^76^ is below that of the resolution of current EM imaging techniques used for connectomic dataset generation. As such, we calculated a proxy for electrical synapse number using the surface area overlap between pairs of neurons. First, we expanded neuron meshes for the haltere afferents and wing-steering MNs by 20nm by displacing all vertices along their own normal (perpendicular to the face). Next, we calculated mesh intersections and the area of this intersection was determined using scikit-image.

#### Motor impact score

The motor impact score^77^ quantifies the net excitatory or inhibitory influence of each haltere afferent subtype on a specific wing-steering MN. It is calculated as a function of the synaptic strength and neurotransmitter identity along all monosynaptic and disynaptic pathways between these neurons. In the fly CNS, acetylcholine is excitatory whilst GABA and glutamate are typically inhibitory^78,79^. As the haltere afferents are cholinergic, we scored monosynaptic connections as the total number of synapses between each haltere afferent subtype and wing-steering MN. To score disynaptic connections, we calculated the fraction of an interneuron’s input that came from each haltere afferent subtype by dividing the number of synapses from that subtype by the interneuron’s total synaptic inputs. We then multiplied this fraction by the number of synapses the interneuron made onto a wing-steering MN to estimate the ‘effective transmitted synapses’ from the haltere afferent to the MN. If the interneuron was cholinergic, the resulting score was considered positive; otherwise, it was considered negative. For each haltere afferent subtype and wing-steering MN, we summed all relevant monosynaptic and disynaptic scores to obtain a final measure of total effective connectivity.

### Design and generation of split-GAL4 lines

To identify enhancers whose expression patterns might label specific haltere afferent subclasses, we used our EM reconstructions to query publicly available GAL4 collections^32,80^ using the color-depth maximum intensity projection (MIP) mask search^81^. In brief, we aligned the EM reconstructions to the JRC2018 VNC template^82^. Next, for each subclass we generated a MIP where color represents z-depth. These MIPs enabled anatomical comparisons against light microscopy images of GAL4 expression patterns through a color-matching algorithm which scores overlapping pixels. We designed putative split-GAL4 combinations for each haltere afferent subclass by choosing two hemidriver lines^83^ whose GAL4 expression patterns were amongst the top ‘hits’ for this subclass. We screened these combinations using flies of the following genotype: pJFRC2-10XUAS-IVS-mCD8::GFP (attp18)/w; Enhancer1-p65ADZp (attP40)/w; Enhancer2-ZpGAL4DBD (attP2)/w. We dissected the VNCs of these animals and imaged native GFP fluorescence. We established correspondence between light microscopy expression patterns and the morphologies of EM reconstructions of haltere afferent subclasses by manual qualitative comparison of key anatomical features. We double balanced hemidriver combinations with strong and consistently reproducible expression patterns and then combined these to make a stable stock.

### Dissection and imaging

#### VNC dissection and immunohistochemistry

We crossed males from each split-GAL4 line to virgin females carrying pJFRC2-10XUAS-IVS-mCD8::GFP in attp18. To ensure consistent GFP expression we chose female progeny aged 5-10 days old for dissection and immunohistochemistry. We anesthetized selected flies with CO2 and briefly submerged them in 70% ethanol before dissecting them in ice-cold PBS as previously described^80^. We fixed isolated VNCs in 4% paraformaldehyde at room temperature for 20 minutes before washing them with PBS-Tx (0.5% Triton X-100 in phosphate-buffered saline). We incubated samples overnight at 4°C with 1:10 mouse anti-nc82 and 1:1000 rabbit anti-GFP blocked with 5% normal goat serum in PBS-Tx. Following primary antibody incubation, we returned samples to room temperature and washed them in PBS-Tx. We applied a secondary antibody stain consisting of 1:250 goat anti-mouse AlexaFluor 633 and 1:250 goat anti-rabbit AlexaFluor 488 overnight at 4°C. Following a final wash in PBS-Tx, we mounted samples in Vectashield antifade reagent (Vector Laboratories, H-1000-10).

#### Haltere dissection

We anesthetized 5-10 day old females with CO2 and briefly submerged them in 70% ethanol. We carefully removed the halteres in 4% paraformaldehyde and then fixed them in this solution for 20 minutes at room temperature. We washed samples in PBS-Tx before mounting them in Vectashield antifade reagent. For all samples, we imaged native fluorescence, i.e. there was no further immunohistochemical amplification.

#### Image acquisition and analysis

We obtained serial optical sections at 0.5μm intervals on a Leica TCS SP8 X using a 40x oil immersion objective at a resolution of 2048 x 2048 pixels. For VNC samples, we set laser excitation to 488 and 633 nm; for haltere samples we took an additional brightfield image. We manually chose the other image acquisition parameters for each sample, with an aim to optimize signal and reduce background noise. These typically consisted of a scan speed of 100 Hz, a line average of 8 and a laser power between 5-10%. Both the VNC and haltere were consistently too large to fit in a single image. Consequently, we captured four tiles per sample and automatically stitched these together following merging and distortion correction using Leica LAS-X software. We carried out further processing of confocal stacks in Fiji (http://fiji.sc/).

## Acknowledgements

This work was supported by NINDS-NIH grant 1U01NS131438-01, a Searle Scholar Award, a McKnight Scholar Award, and NSF grant IOS2221458 to B.H.D. The content is solely the responsibility of the authors and does not necessarily represent the official views of the National Institutes of Health. We thank J. Fox, M. Murthy, and J. Tuthill for comments on the manuscript.

## Author contributions

S.D. and B.H.D conceived the project; B.H.D. acquired funding. S.D. proofread neurons in FANC. S.D. and Z.H. performed light microscopic imaging. S.D. analyzed the data. S.D. and B.H.D. wrote the paper.

## Extended Data figure legends

**Extended Data Figure 1:**
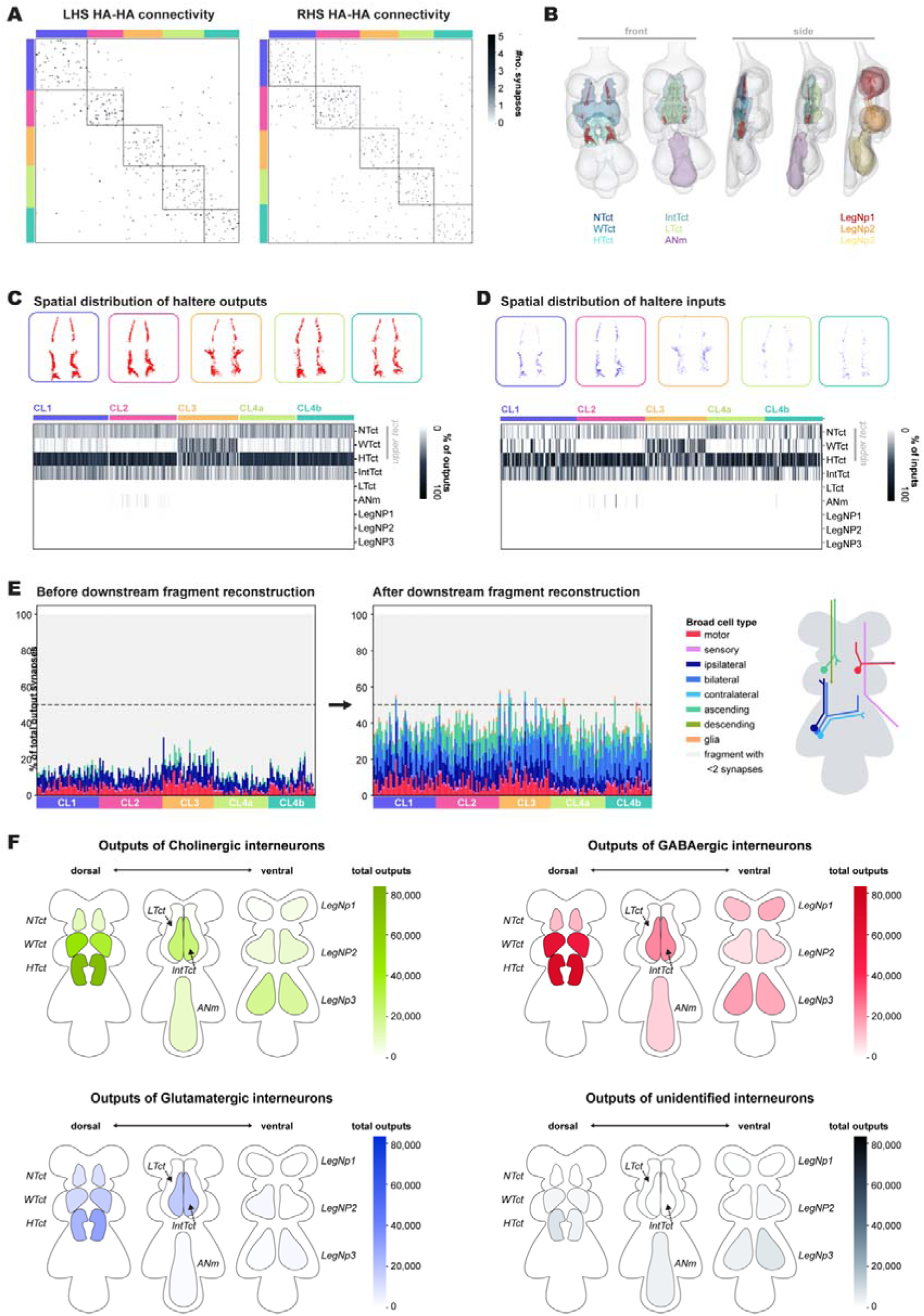
Quantifying and localizing the synaptic inputs and outputs of the haltere afferents and its interneurons. A: Heatmaps showing axo-axonic synapses between the haltere afferents. B: Rendering of the haltere synaptic outputs (red dots) within different VNC neuropils. C: Rendering of the synaptic outputs for each haltere afferent subtype, alongside a heatmap showing the normalized distribution of these outputs within different VNC neuropils for all individual neurons. D: Same as A but for synaptic inputs. E: Stacked barcharts showing the percentage of outputs from each LHS haltere afferent to different broad morphological classes of neuron before (top) and after (bottom) fragment reconstruction. F: Schematized heatmaps depicting the spatial distribution of the synaptic outputs of interneurons directly downstream of the haltere afferents within different neuropils of the VNC. Interneurons are grouped by putative neurotransmitter type as determined by hemilineage-typing.

**Extended Data Figure 2:**
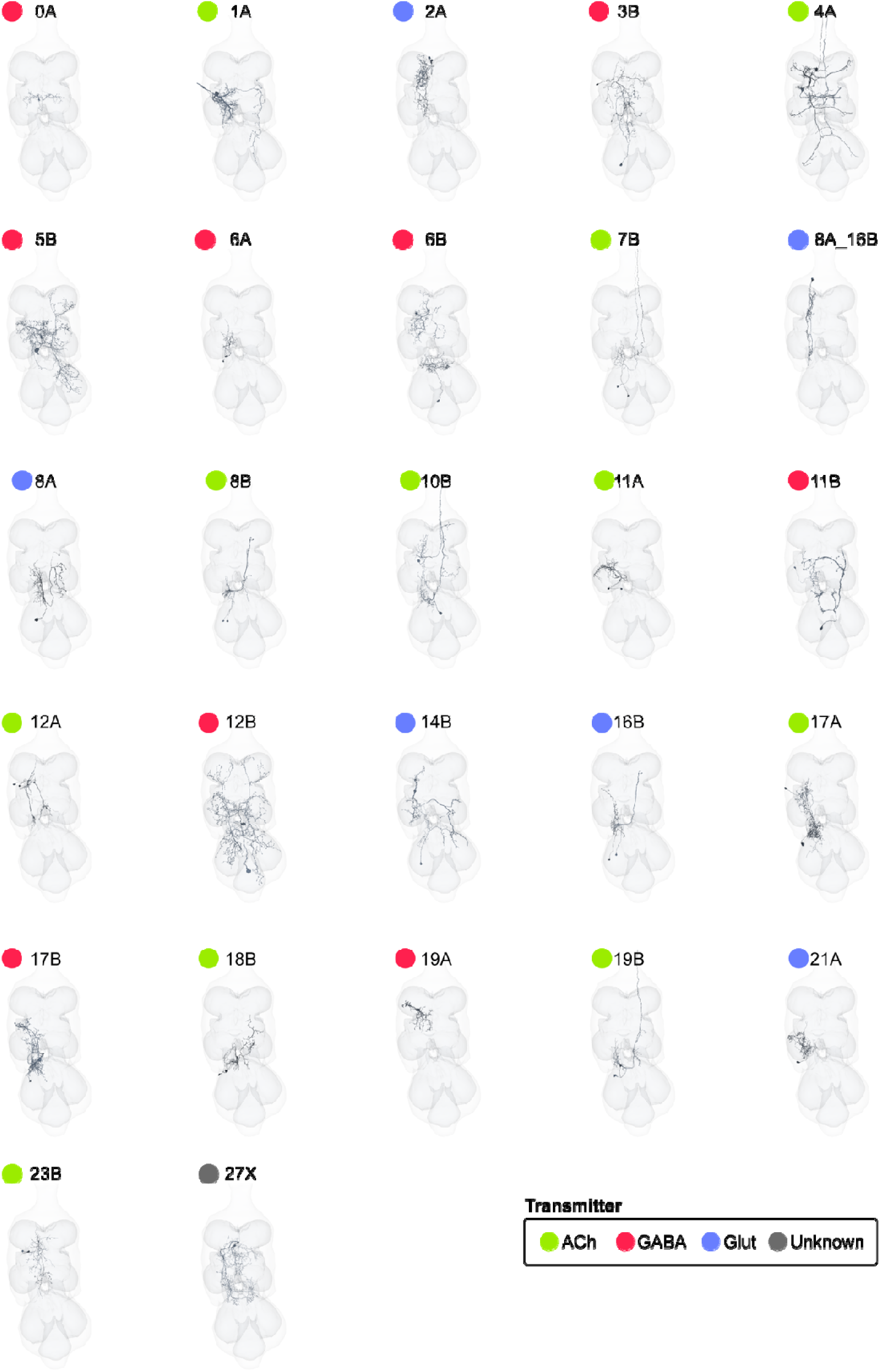
Interneurons that receive haltere input. Example interneurons from each hemilineage. Colors depict neurotransmitter identity (acetylcholine = green, GABA = red, glutamate = blue).

**Extended Data Figure 3:**
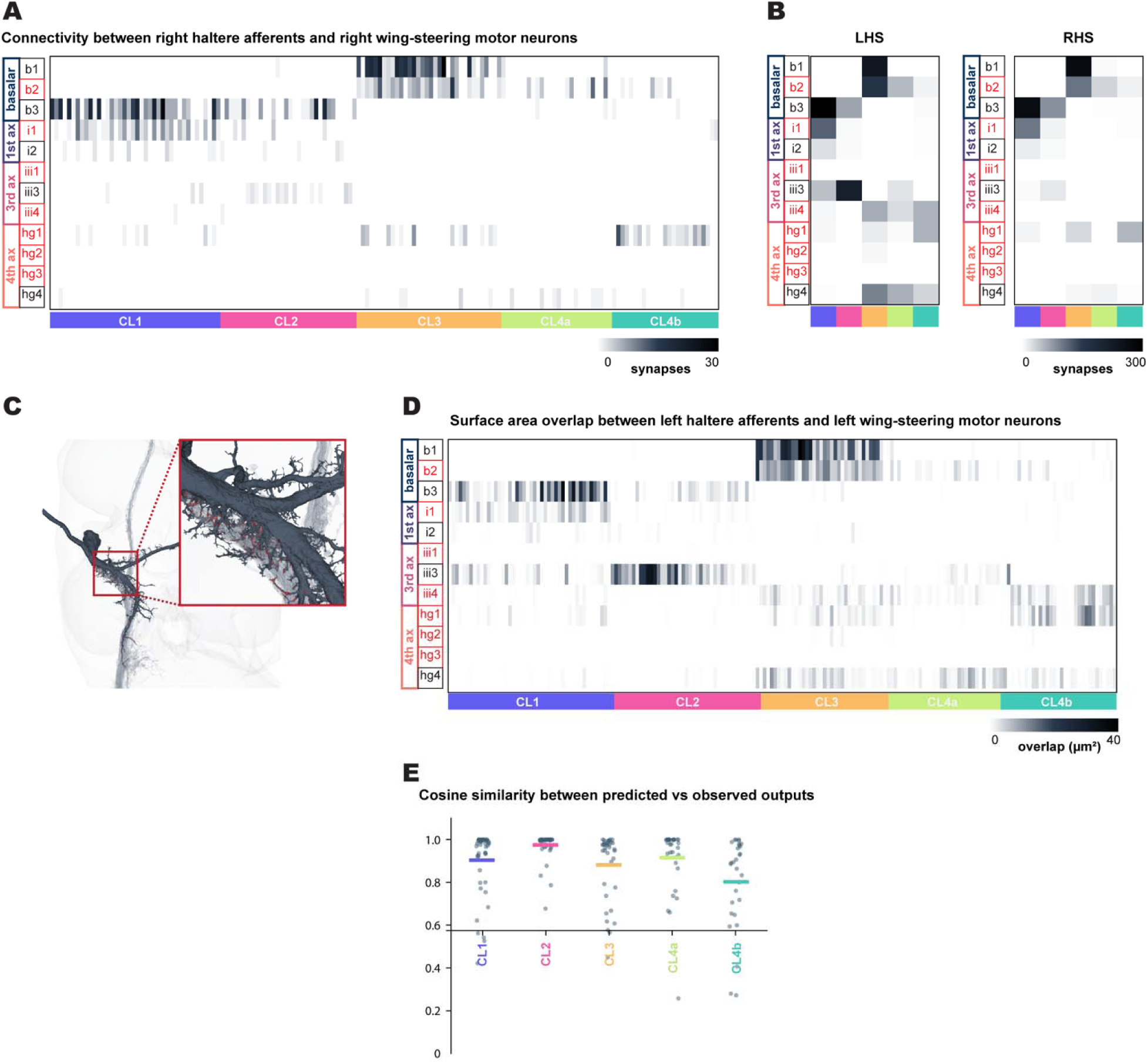
Quantification of putative electrical synapses between the haltere afferents and wing steering muscle motor neurons. A: Connectivity between the RHS haltere sensory afferents and the wing steering muscle motor neurons. Rows are grouped by the sclerite which a wing-steering muscle attaches to and colored by functional classification: tonic (black) or phasic (red); columns are colored by haltere morphological subtype. B: Bilateral comparison of the motor impact of each afferent morphological cluster on a given wing steering muscle motor neuron. Each comparison is between haltere afferents and ipsilateral wing steering muscle motor neurons. C: Example of surface area contact between haltere cluster 3 and the b1 wing-steering motor neuron. Intersection area is highlighted in red. D: Mean surface area contact between the haltere afferents and wing-steering motor neurons. E: Cosine similarity of vectors for predicted electrical synapses between the haltere afferents and the wing-steering motor neurons, as determined by surface area overlap, and the observed chemical synaptic connectivity in the FANC dataset.

**Extended Data Figure 4:**
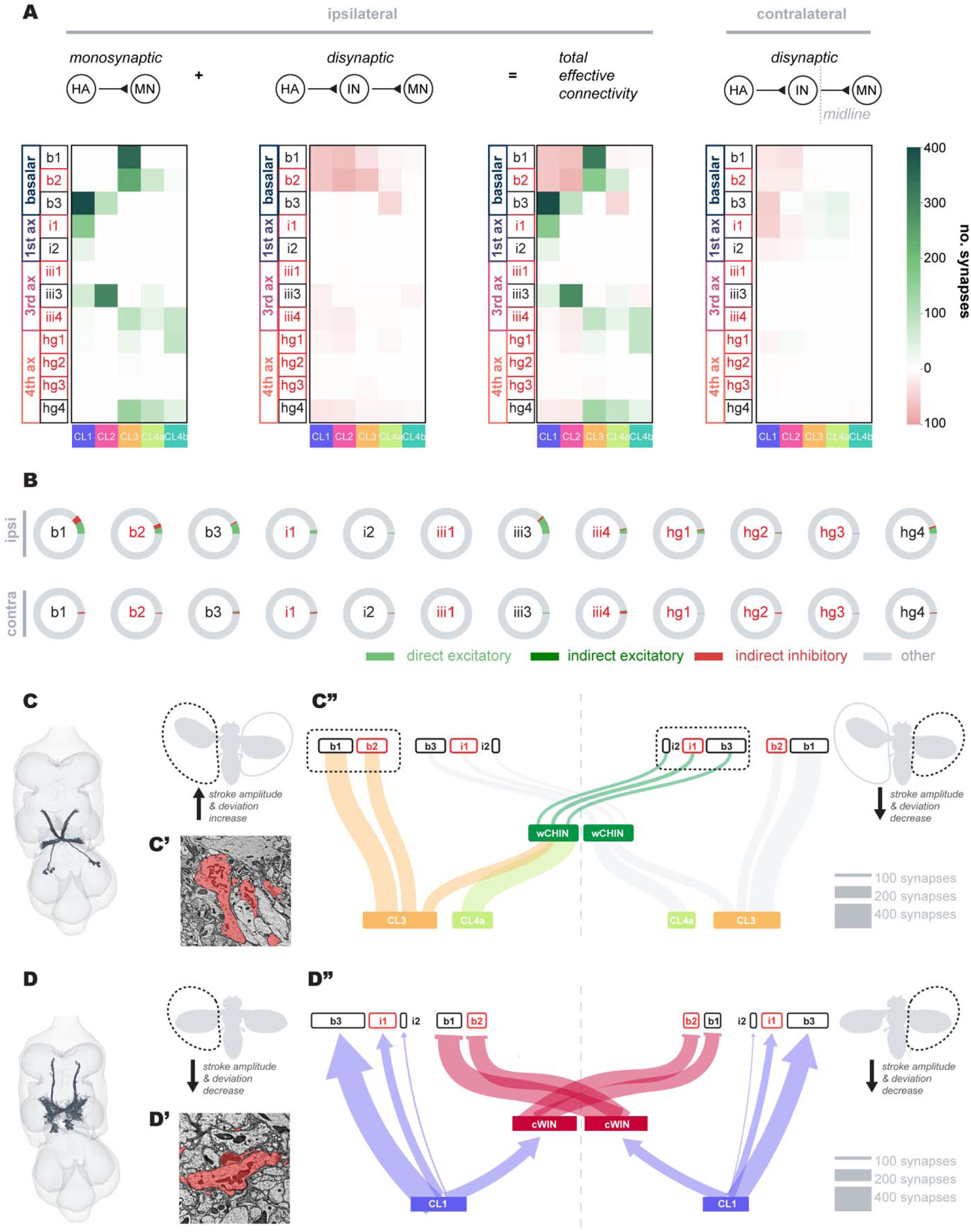
Bilateral coordination of wing kinematics. A: The first three heatmaps depict the direct (monosynaptic) and indirect (disynaptic) connections between the LHS haltere afferents and the LHS wing-steering MNs, as well as the total effective connectivity between these groups (summed monosynaptic and disynaptic connectivity). The fourth heatmap depicts the indirect connectivity between the LHS haltere afferents and the RHS wing-steering MNs. B: Donut charts illustrating the percentage of synaptic input to each wing-steering MN received as direct excitatory connections from the LHS afferents vs indirect excitatory/inhibitory connections via a single interneuron. C: Rendering of the contralateral haltere interneurons (cHINs). C’: Exemplary EM image showing densely packed mitochondria at the axon terminals of a cHIN, which might be indicative of electrotonic transmission. C’’: Summary wiring diagram of connectivity between the haltere afferents, cHINs and wing-steering motor neurons. Haltere subclasses that ipsilaterally activate wing-steering motor neurons that increase wingstroke amplitude can contralaterally activate wing-steering motor neurons that decrease wingstroke amplitude via the cHIN interneurons. This allows for asymmetric steering maneuvers. D: Same as C, but for contralateral wing interneurons (cWINs). In contrast to the cHINs, haltere subclasses that help decreases in wingstroke amplitude may contralaterally activate wing steering muscle motor neurons that decrease wingstroke amplitude.

## References

1. Knudsen, E. I., S Lac, A. & Esterly, S. D. Computational Maps in the Brain. (2003) doi:10.1146/annurev.ne.10.030187.000353.

2. Carr, C. E. Processing of temporal information in the brain. Annu. Rev. Neurosci. 16, 223– 243 (1993).

3. Tu, M. S. & Dickinson, M. H. The control of wing kinematics by two steering muscles of the blowfly (Calliphora vicina). J Comp Physiol A 178, 813–830 (1996).

4. Heide, G. & Götz, K. G. Optomotor control of course and altitude in Drosophila melanogaster is correlated with distinct activities of at least three pairs of flight steering muscles. J Exp Biol 199, 1711–1726 (1996).

5. Balint, C. N. & Dickinson, M. H. The correlation between wing kinematics and steering muscle activity in the blowfly Calliphora vicina. J. Exp. Biol. 204, 4213–4226 (2001).

6. Lindsay, T., Sustar, A. & Dickinson, M. The Function and Organization of the Motor System Controlling Flight Maneuvers in Flies. Curr Biol 27, 345–358 (2017).

7. Heide, G. Neural mechanisms of flight control in Diptera. BIONA-Rep. 2, 35–52 (1983).

8. Azevedo, A. et al. Connectomic reconstruction of a female Drosophila ventral nerve cord. Nature 631, 360–368 (2024).

9. Phelps, J. S. et al. Reconstruction of motor control circuits in adult Drosophila using automated transmission electron microscopy. Cell 184, 759–774.e18 (2021).

10. Pringle, J. W. S. The gyroscopic mechanism of the halteres of Diptera. Philos Trans R Soc Lond B Biol Sci 233, 347–384 (1997).

11. Fraenkel, G. & Pringle, J. W. S. Biological Sciences: Halteres of Flies as Gyroscopic Organs of Equilibrium. Nature 141, 919–920 (1938).

12. Cole, E. S. & Palka, J. The pattern of campaniform sensilla on the wing and haltere of Drosophila melanogaster and several of its homeotic mutants. Development 71, 41–61 (1982).

13. Derham, W. Physico-Theology: Or, a Demonstration of the Being and Attributes of G Od, from His Works of Creation. (Robinson and Roberts, 1768).

14. Nalbach, G. The halteres of the blowfly Calliphora. I. Kinematics and dynamics. J. Comp. Physiol. A 173, 293–300 (1993).

15. Uga, S. & Kuwabara, M. The fine structure of the campaniform sensilium on the haltere of the fleshfly, boettcherisca peregrina. J. Electron Microsc. (Tokyo) 16, 304–312 (1967).

16. Smith, D. S. The fine structure of haltere sensilla in the blowfly Calliphora erythrocephala (Meig.), with scanning electron microscopic observations on the haltere surface. Tissue Cell 1, 443–484 (1969).

17. Toh, Y. Structure of campaniform sensilla on the haltere ofDrosophila prepared by cryofixation. J. Ultrastruct. Res. 93, 92–100 (1985).

18. Chan, W. P. & Dickinson, M. H. Position-specific central projections of mechanosensory neurons on the haltere of the blow fly, Calliphora vicina. J Comp Neurol 369, 405–418 (1996).

19. Agrawal, S., Grimaldi, D. & Fox, J. L. Haltere morphology and campaniform sensilla arrangement across Diptera. Arthropod Struct Dev 46, 215–229 (2017).

20. Hengstenberg, R. Mechanosensory control of compensatory head roll during flight in the blowfly Calliphora erythrocephala Meig. J. Comp. Physiol. A 163, 151–165 (1988).

21. Dickinson, M. H. Haltere-mediated equilibrium reflexes of the fruit fly, Drosophila melanogaster. Philos Trans R Soc Lond B Biol Sci 354, 903–916 (1999).

22. Sherman, A. & Dickinson, M. H. A comparison of visual and haltere-mediated equilibrium reflexes in the fruit fly Drosophila melanogaster. J. Exp. Biol. 206, 295–302 (2003).

23. Sherman, A. & Dickinson, M. H. Summation of visual and mechanosensory feedback in Drosophila flight control. J. Exp. Biol. 207, 133–142 (2004).

24. Bender, J. A. & Dickinson, M. H. A comparison of visual and haltere-mediated feedback in the control of body saccades in Drosophila melanogaster. J Exp Biol 209, 4597–4606 (2006).

25. Huston, S. J. & Krapp, H. G. Nonlinear integration of visual and haltere inputs in fly neck motor neurons. J Neurosci 29, 13097–13105 (2009).

26. Trimarchi, J. R. & Murphey, R. K. The shaking-B2 mutation disrupts electrical synapses in a flight circuit in adult Drosophila. J Neurosci 17, 4700–4710 (1997).

27. Fayyazuddin, A. & Dickinson, M. H. Haltere afferents provide direct, electrotonic input to a steering motor neuron in the blowfly, Calliphora. J Neurosci 16, 5225–5232 (1996).

28. Court, R. et al. A systematic nomenclature for the Drosophila ventral nerve cord. Neuron 107, 1071–1079.e2 (2020).

29. Dickerson, B. H., de Souza, A. M., Huda, A. & Dickinson, M. H. Flies Regulate Wing Motion via Active Control of a Dual-Function Gyroscope. Curr Biol 29, 3517–3524.e3 (2019).

30. Costa, M., Manton, J. D., Ostrovsky, A. D., Prohaska, S. & Jefferis, G. S. X. E. NBLAST: Rapid, Sensitive Comparison of Neuronal Structure and Construction of Neuron Family Databases. Neuron 91, 293–311 (2016).

31. Dickerson, B. H., Fox, J. L. & Sponberg, S. Functional diversity from generic encoding in insect campaniform sensilla. Curr. Opin. Physiol. 19, 194–203 (2021).

32. Meissner, G. W. et al. A searchable image resource of Drosophila GAL4 driver expression patterns with single neuron resolution. Elife 12, (2023).

33. Shepherd, G. M., Rowe, T. B. & Greer, C. A. An evolutionary microcircuit approach to the neural basis of high dimensional sensory processing in olfaction. Front Cell Neurosci 15, 658480 (2021).

34. Kazama, H. & Wilson, R. I. Origins of correlated activity in an olfactory circuit. Nat Neurosci 12, 1136–1144 (2009).

35. Cover, K. K. & Mathur, B. N. Axo-axonic synapses: Diversity in neural circuit function. J Comp Neurol 529, 2391–2401 (2021).

36. Gruber, L. et al. The unique synaptic circuitry of specialized olfactory glomeruli in Drosophila melanogaster. eLife (2023) doi:10.7554/elife.88824.1.

37. Namiki, S., Dickinson, M. H., Wong, A. M., Korff, W. & Card, G. M. The functional organization of descending sensory-motor pathways in Drosophila. Elife 7, (2018).

38. Lesser, E. et al. Synaptic architecture of leg and wing premotor control networks in Drosophila. Nature 631, 369–377 (2024).

39. Lacin, H. et al. Neurotransmitter identity is acquired in a lineage-restricted manner in the Drosophila CNS. Elife 8, e43701 (2019).

40. Harris, R. M., Pfeiffer, B. D., Rubin, G. M. & Truman, J. W. Neuron hemilineages provide the functional ground plan for the Drosophila ventral nervous system. Elife 4, (2015).

41. Boettiger, E. G. & Furshpan, E. The mechanics of flight movements in diptera. Biol. Bull. 102, 200–211 (1952).

42. Miyan J. A., Ewing A. W., & Lighthill Michael James. How Diptera move their wings: a re-examination of the wing base articulation and muscle systems concerned with flight. Philos. Trans. R. Soc. Lond. B Biol. Al Sci. 311, 271–302 (1985).

43. Dickinson, M. H. & Tu, M. S. The function of dipteran flight muscle. Comp. Biochem. Physiol. A Physiol. 116, 223–238 (1997).

44. Bhandawat, V., Olsen, S. R., Gouwens, N. W., Schlief, M. L. & Wilson, R. I. Sensory processing in the Drosophila antennal lobe increases reliability and separability of ensemble odor representations. Nat Neurosci 10, 1474–1482 (2007).

45. Verbe, A., Lea, K. M., Fox, J. L. & Dickerson, B. H. Flies tune the activity of their multifunctional gyroscope. Curr. Biol. 34, 3644–3653.e3 (2024).

46. Dickerson, B. H. Timing precision in fly flight control: integrating mechanosensory input with muscle physiology. Proc Biol Sci 287, 20201774 (2020).

47. Chan, W. P., Prete, F. & Dickinson, M. H. Visual input to the efferent control system of a fly’s ‘gyroscope’. Science 280, 289–292 (1998).

48. Chapman, K. M. & Smith, R. S. A linear transfer function underlying impulse frequency modulation in a cockroach mechanoreceptor. Nature 197, 699 (1963).

49. Melis, J. M., Siwanowicz, I. & Dickinson, M. H. Machine learning reveals the control mechanics of an insect wing hinge. Nature 628, 795–803 (2024).

50. Trimarchi, J. R. & Schneiderman, A. M. The motor neurons innervating the direct flight muscles of Drosophila melanogaster are morphologically specialized. J. Comp. Neurol. 340, 427–443 (1994).

51. Fayyazuddin, A. & Dickinson, M. H. Convergent mechanosensory input structures the firing phase of a steering motor neuron in the blowfly, Calliphora. J Neurophysiol 82, 1916–1926 (1999).

52. Tu, M. & Dickinson, M. Modulation of negative work output from a steering muscle of the blowfly Calliphora Vicina. J Exp Biol 192, 207–224 (1994).

53. Lehmann, F. O. & Götz, K. G. Activation phase ensures kinematic efficacy in flight-steering muscles of Drosophila melanogaster. J Comp Physiol A 179, 311–322 (1996).

54. Strausfeld, N. J. & Seyan, H. S. Convergence of visual, haltere, and prosternal inputs at neck motor neurons of Calliphora erythrocephala. Cell Tissue Res 240, 601–615 (1985).

55. Oldfield, B. P. Tonotopic organisation of auditory receptors in tettigoniidae (Orthoptera: Ensifera). J. Comp. Physiol. 147, 461–469 (1982).

56. Field, L. H. Mechanism for range fractionation in chordotonal organs of Locusta migratoria (L) and Valanga sp. (Orthoptera : Acrididae). Int. J. Insect Morphol. Embryol. 20, 25–39 (1991).

57. Matheson, T. Range fractionation in the locust metathoracic femoral chordotonal organ. J. Comp. Physiol. A 170, 509–520 (1992).

58. Fettiplace, R. & Fuchs, P. A. Mechanisms of hair cell tuning. Annu. Rev. Physiol. 61, 809– 834 (1999).

59. Dickinson, M. H. Linear and nonlinear encoding properties of an identified mechanoreceptor on the fly wing measured with mechanical noise stimuli. J. Exp. Biol. 151, 219–244 (1990).

60. Dickinson, M. H. Comparison of encoding properties of campaniform sensilla on the fly wing. J. Exp. Biol. 151, 245–261 (1990).

61. Fox, J. L., Fairhall, A. L. & Daniel, T. L. Encoding properties of haltere neurons enable motion feature detection in a biological gyroscope. Proc Natl Acad Sci U A 107, 3840–3845 (2010).

62. Yarger, A. M. & Fox, J. L. Single mechanosensory neurons encode lateral displacements using precise spike timing and thresholds. Proc Biol Sci 285, (2018).

63. Jacobs, G. A. & Theunissen, F. E. Functional organization of a neural map in the cricket cercal sensory system. J Neurosci 16, 769–784 (1996).

64. Paydar, S., Doan, C. A. & Jacobs, G. A. Neural mapping of direction and frequency in the cricket cercal sensory system. J. Neurosci. 19, 1771–1781 (1999).

65. Knudsen, E. I. & Konishi, M. A neural map of auditory space in the owl. Science 200, 795– 797 (1978).

66. Carr, C. E. & Konishi, M. A circuit for detection of interaural time differences in the brain stem of the barn owl. J Neurosci 10, 3227–3246 (1990).

67. Palka, J., Malone, M., Ellison, R. & Wigston, D. Central projections of identified Drosophila sensory neurons in relation to their time of development. J Neurosci 6, 1822–1830 (1986).

68. Dickinson, M. H. & Palka, J. Physiological properties, time of development, and central projection are correlated in the wing mechanoreceptors of Drosophila. J Neurosci 7, 4201– 4208 (1987).

69. Lesser, E., A Moussa & Tuthill, J. Peripheral anatomy and central connectivity of proprioceptive sensory neurons in the Drosophila wing. bioRxiv (2025).

70. Takemura, S.-Y., et al. A Connectome of the Male Drosophila Ventral Nerve Cord. bioRxiv (2023) doi:10.1101/2023.06.05.543757.

71. Grimaldi, D. & Engel, M. S. Evolution of the Insects. (Cambridge University Press, 2005).

72. Maitin-Shepard, J., et al. google/neuroglancer: 10.5281/zenodo.5573294 (2021).

73. Stürner, T. et al. Comparative connectomics of the descending and ascending neurons of the Drosophila nervous system: stereotypy and sexual dimorphism. bioRxivorg 2024.06.04.596633 (2024) doi:10.1101/2024.06.04.596633.

74. Dorkenwald, S. et al. CAVE: Connectome Annotation Versioning Engine. Nat. Methods 22, 1112–1120 (2025).

75. Shepherd, D., Harris, R., Williams, D. W. & Truman, J. W. Postembryonic lineages of the Drosophila ventral nervous system: Neuroglian expression reveals the adult hemilineage associated fiber tracts in the adult thoracic neuromeres. J Comp Neurol 524, 2677–2695 (2016).

76. Ammer, G., Vieira, R. M., Fendl, S. & Borst, A. Anatomical distribution and functional roles of electrical synapses in Drosophila. Curr Biol 32, 2022–2036.e4 (2022).

77. Lee, S.-Y. J., Dallmann, C. J., Cook, A., Tuthill, J. C. & Agrawal, S. Divergent neural circuits for proprioceptive and exteroceptive sensing of the Drosophila leg. 2024.04.23.590808 Preprint at 10.1101/2024.04.23.590808 (2024).

78. Allen, A. M. et al. A single-cell transcriptomic atlas of the adult Drosophila ventral nerve cord. eLife 9, e54074 (2020).

79. Liu, W. W. & Wilson, R. I. Glutamate is an inhibitory neurotransmitter in the Drosophila olfactory system. Proc. Natl. Acad. Sci. U. S. A. 110, 10294–10299 (2013).

80. Jenett, A. et al. A GAL4-driver line resource for Drosophila neurobiology. Cell Rep 2, 991– 1001 (2012).

81. Otsuna, H., Ito, M. & Kawase, T. Color depth MIP mask search: a new tool to expedite Split-GAL4 creation. *bioRxiv* (2018) doi:10.1101/318006.

82. Bogovic, J. A. et al. An unbiased template of the Drosophila brain and ventral nerve cord. PLoS One 15, e0236495 (2020).

83. Dionne, H., Hibbard, K. L., Cavallaro, A., Kao, J.-C. & Rubin, G. M. Genetic Reagents for Making Split-GAL4 Lines in Drosophila. Genetics 209, 31–35 (2018).

